# VirPreNet: A weighted ensemble convolutional neural network for the virulence prediction of influenza A virus using all 8 segments

**DOI:** 10.1101/2020.07.31.230904

**Authors:** Rui Yin, Zihan Luo, Pei Zhuang, Zhuoyi Lin, Chee Keong Kwoh

## Abstract

**Motivation:** Influenza viruses are persistently threatening public health, causing annual epidemics and sporadic pandemics. The evolution of influenza viruses remains to be the main obstacle in the effectiveness of antiviral treatments due to rapid mutations. Previous work has been investigated to reveal the determinants of virulence of the influenza A virus. To further facilitate flu surveillance, explicit detection of influenza virulence is crucial to protect public health from potential future pandemics.

**Results:** In this paper, we propose a weighted ensemble convolutional neural network for the virulence prediction of influenza A viruses named VirPreNet that uses all 8 segments. Firstly, mouse lethal dose 50 is exerted to label the virulence of infections into two classes, namely avirulent and virulent. A numerical representation of amino acids named ProtVec is applied to the 8-segments in a distributed manner to encode the biological sequences. After splittings and embeddings of influenza strains, the ensemble convolutional neural network is constructed as the base model on the influenza dataset of each segment, which serves as the VirPreNet’s main part. Followed by a linear layer, the initial predictive outcomes are integrated and assigned with different weights for the final prediction. The experimental results on the collected influenza dataset indicate that VirPreNet achieves state-of-the-art performance combining ProtVec with our proposed architecture. It outperforms baseline methods on the independent testing data. Moreover, our proposed model reveals the importance of PB2 and HA segments on the virulence prediction. We believe that our model may provide new insights into the investigation of influenza virulence.

**Availability and Implementation:** Codes and data to generate the VirPreNet are publicly available at https://github.com/Rayin-saber/VirPreNet

## 1 Introduction

Influenza A virus can easily infect the respiratory and cause a highly contagious disease. It consists of 8 single-stranded, negative-sense viral RNA segments that encode at least 12 different proteins (Bouvier and Palese, 2008). The viruses possess mutability and high frequency of genetic reassortment (Vijaykrishna et al., 2015; Cheung et al., 2015; Yin et al., 2020a). Hence, they will have a great ability to be virulent and cause high mortality and morbidity when infecting humans. In addition to seasonal influenza epidemics, when millions of people are infected worldwide and up to 500,000 people are killed every year according to the World Health Organization (WHO) (Organization et al., 2009), influenza viruses are responsible for severe outbreaks in history, e.g., the 1918 H1N1 Spanish Flu, the 1957 H2N2 Asian Flu, the 1968 H3N2 Hong Kong Flu, the 1977 H1N1 Russia Flu and the most recent 2009 Swine-origin Flu (Saunders-Hastings and Krewski, 2016). The pandemic strains are still circulating among humans and continuously lead to recurrent epidemics. It is shown that several other subtypes have also infected humans, including H5N1, H5N6, H6N1, H7N2, H7N7, H7N9, H9N2 and H10N8 (Poovorawan et al., 2013; Su et al., 2015). Among them, H5N1 and H7N9 have raised a major public concern due to their ability to infect humans with a high fatality rate (Ma et al., 2018). Overall, the detection of virulence level is critical to estimate the lethality of influenza viruses and facilitate flu surveillance for better precautions.

Current systems of flu surveillance by WHO rely on empirical determination of antigenicity through biological experiments such as hemagglutination inhibition (HI) assays (Ndifon et al., 2009) and micro-neutralization assays (Wu et al., 2016). However, these methods are time-consuming and labour-intensive to determine the antigenicity of influenza viruses. Virulence measures the degree of pathogenicity and can reflect the capability of the virus to cause disease. It is a more accurate correlation to the viral lethality compared with antigenicity and provides insights for a potential epidemic or pandemic. Mutations and reassortment in the influenza viruses will cause antigenic drift and shift that makes the protein unrecognizable to pre-existing host immunity (Zhou et al., 2018; Yin et al., 2020b). Previous studies mainly focused on the variations in viral genes that influence virulence. For example, the mutations in the region 130-loop, 190-helix and 220-loop during the adaptation of influenza A virus in mice, which surround the receptor-binding site in the HA protein, have increased its virulence (Imai and Kawaoka, 2012). The E627K and D701N in protein PB2 have also been considered as biological markers for the virulence of influenza viruses in mice (Kamal et al., 2014). In addition, Yin et al. identified several critical virulent sites that are associated with past pandemic strains (Yin et al., 2017). Sander at al. selected mutant A/H5N1 viruses, indicating Q222L and G224S changed the receptor binding of H2 and H3 avian influenza binding specificity to alpha (2,6) linked sialic acid, which contributed to the outbreak of 1957 and 1968 pandemics (Zhang et al., 2013). The dual mutation S224P and N383D in protein PA caused the increase of polymerase activity and has been regarded as a hallmark for natural adaptation of H1N1 and H5N1 viruses to mammals (Song et al., 2015). Chinh et al. found Mutation R289K-induced conformational in H7N9 suggests potential adaptions of the virus itself for future drug-resistance (Su et al., 2013). Apart from the mutations in proteins HA, PB2 and PA, amino acid substitutions at 223 and 275 in NA protein (de Vries et al., 2012; LeGoff et al., 2012) and 92 in NS1 protein (Seo et al., 2002) have also made an effect with enhanced virulence in mammalian hosts (Yamada et al., 2010). These studies demonstrated that the amino acid substitutions on different viral proteins could make an impact on the degree of virulence.

With increasingly available biological sequences and advancement of computing capacity, it is highly desired to develop computational methods for rapidly and effectively detecting viral virulence or its associated factors. The following list is by no means exhaustive but a small sample of previous approaches that explore virulence detection. Garg and Gupta illustrated a bacterial virulent protein prediction method through a bi-layer cascade support vector machine (Garg and Gupta, 2008). Zheng et al. showed a novel network-based for identifying the virulence factors in the proteomes based on the protein-protein interaction networks (Zheng et al., 2012). Moreover, Ivan et al. developed a small benchmark dataset for the influenza A virus and explored rule-based approaches for classifying the virulent strains (Ivan and Kwoh, 2019). Of these approaches, it has demonstrated the feasibility of utilizing computational methods in exploring viral virulence. The recent development of deep learning enables enormous progress of predictive tasks in various fields.

In this article, we propose a weighted ensemble CNN named VirPreNet that incorporates 8 different influenza segments to predict the virulence of influenza A viruses. Specifically, we introduce ProtVec (Asgari and Mofrad, 2015), a continuous distributed representation of amino acids, that encodes influenza strains into numerical vectors. We combine this vector-based space representation with a deep learning architecture, i.e. ResNeXt (Xie et al., 2017) as the base model, which adopts the strategy of repeating layers of ResNet for the virulence prediction on the single-segmented dataset. The base model trained on the data of each segment is integrated through a linear layer for the final prediction. It is shown that VirPreNet can not only adapt to a variety of networks independently but also learn different weights for the importance of base model on distinct influenza segments. We conduct experiments on the collected influenza dataset. It is empirically indicated that VirPreNet can improve the prediction accuracy using all 8 segments. Compared with the baseline methods, our proposed model achieves better or comparable performance on virulence prediction. To the best of our knowledge, this is the first time that attempts to apply deep learning techniques for virulence prediction using ensemble CNN. It also assigns the learned weights on base models, outlining the importance of different proteins of influenza viruses. The encouraging predictive results suggest that our method can be potentially used for flu surveillance to estimate the lethality of the novel emerging influenza strains.

## 2 Materials and Methods

### 2.1 Definition of virulence

The definition of virulence is generally regarded as the capability of a pathogen or microbe to infect or damage a host (Thrall and Burdon, 2003). More specifically in animal systems, it refers to the degree of damage caused by a microbe to its host (Pirofski and Casadevall, 2012). In this work, we only focus on the virulence of the influenza A virus. As far as we know, virulence has not been clearly defined with a rigorous mathematical definition. Here we leverage the mouse lethal dose (MLD) 50 (da Costa et al., 2015), the time series of weight loss or percentage of survival, to measure the virulence of infections. Hence, the degree of virulence is categorized into two classes, namely avirulent and virulent, which are labeled as “0” and “1”, respectively. If the MLD50 is greater than 10E6.0 (regardless of its unit), we consider it as avirulent. Otherwise, it is regarded as virulent. Besides, if the virulence level of infection cannot be determined from the upper or lower bound of MLD50, we follow RULE 1 to 6 in (Ivan and Kwoh, 2019) to classify the remaining samples.

### 2.2 Data collection and processing

The datasets contain sequence data and virulence information of influenza A virus infections. We collect virulence information from previous publications and experiments (Ivan and Kwoh, 2019). The MLD 50 information is recorded in each infection with specific influenza A virus strain and mouse strain. If the information that does not contains MLD50 value, but the time series of weight loss or percentage of survival of infected mice per infection dose could be inferred from the relevant figures, the lower or upper bound of MLD50 values are also used to estimate the virulence label. The sequence data in the literature information consisted of all segments downloaded from Influenza Virus Resource (IVR) (Sayers et al., 2012). If a genomic segment of a particular virus is incomplete, basic local alignment search tool (BLAST) (Altschul et al., 1990) would be applied to search the top virus whose genomes are available to extrapolate the incomplete genome. Besides, the reassortant strains are reconstructed using relevant viral segments. Finally, it is ended up with 489 records of influenza A virus information with corresponding complete genome strains.

To process the collected influenza strains from different subtypes, multiple alignment using fast Fourier transform (MAFFT) (Katoh and Standley, 2013)) is performed for each segment along with their target virulence class. These sequences are comprised of the alignments of all influenza proteins. The H3 and N2 numbering system (Burke and Smith, 2014) is utilized to label the position in the alignments of HA and NA, respectively. Apart from the 20 common amino acids that are encoded by the codons of the universal genetic code directly, there exist several ambiguous amino acids in some of the strains. These amino acids where chemical or crystallographic analysis of a peptide or protein can not conclusively determine the identity of a residue (Thrall and Burdon, 2003). They are also adopted to summarize conserved protein sequence motifs. The abbreviation of amino acids to indicate the sets of similar residues contains ‘B’ (‘D’, ‘N’), ‘Z’ (‘E’, ‘Q’), ‘J’ (‘I’, ‘L’) and ‘X’ (all amino acids), where the letters in the bracket are the potential amino acids representing the ambiguous residues. We randomly replace these four ambiguous amino acids with substitutions by the biological embedding for the feature space construction.

### 2.3 The VirPreNet framework

Fig. 1 shows an overview of VirPreNet. Briefly, we first split the raw sequence proteins into subsequences with overlapped 3-grams, which are embedded based on ProtVec (Asgari and Mofrad, 2015), a continuous distributed representation of biological sequences. The embedded vectors of each influenza segment are respectively fed into base model ResNeXt that aggregates a set of transformations with the same topology to independently predict virulence label. A linear layer is added that incorporates all the predictive results from the previous layer for the final prediction. The details of each step will be described in the following subsections.

**Fig. 1.**
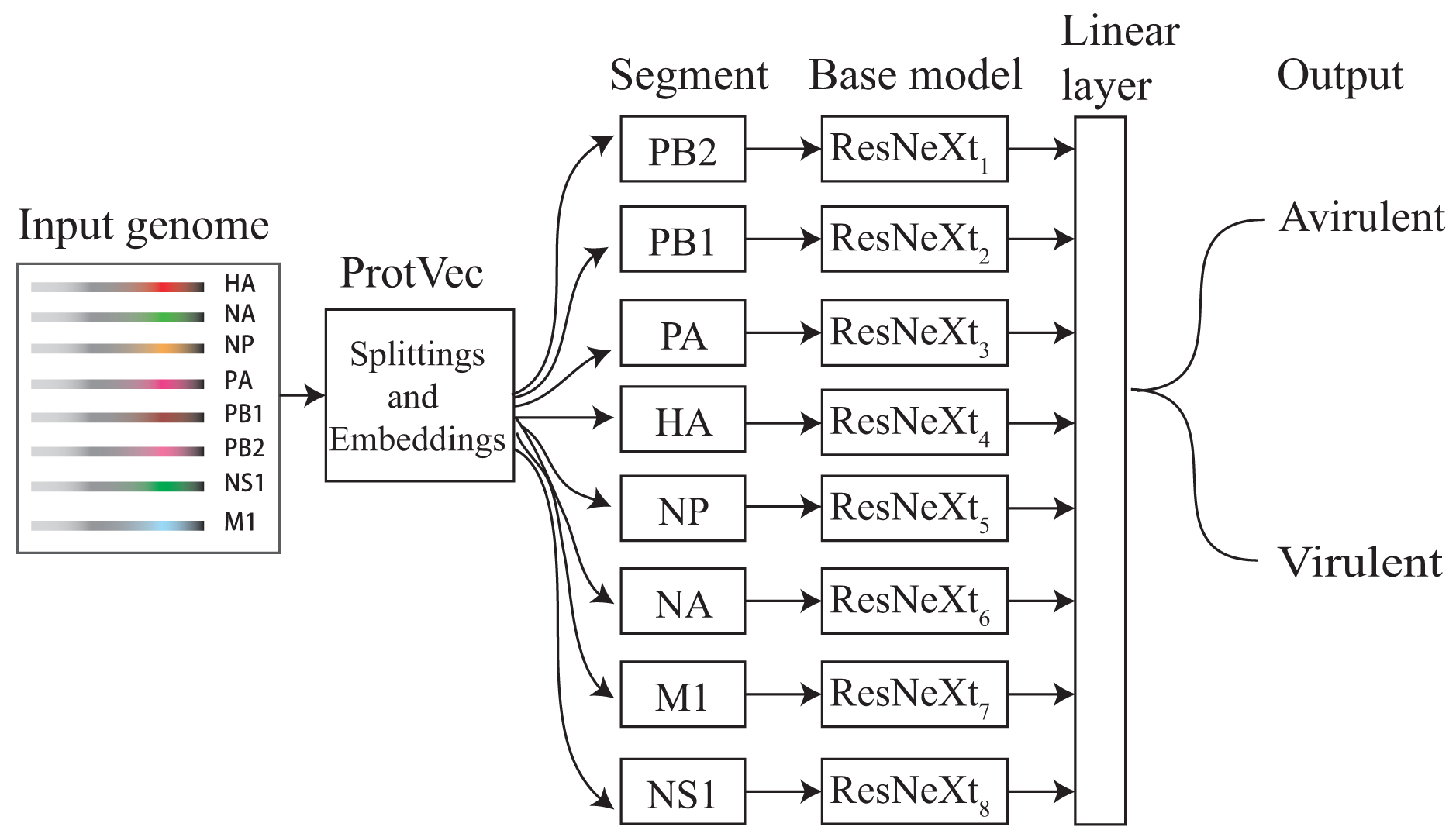
The overview of the proposed framework VirPreNet. The input genome data is split and embedded based on ProtVec. The influenza data of each segment is fed to the base ResNeXt model. Each head outputs the outcome of each class is the correct one, followed by a linear layer to determine the final label.

#### 2.3.1 Feature space

The introduction of distributed representation has been successfully applied in natural language processing by mapping the words or sentences into real-value vectors. In bioinformatics tasks, it is critical to express the biological information with a discrete model or a vector because sequence data is not directly applicable to the existing machine learning approaches (Chou, 2015). A recent distributed representation ProtVec has been proposed for bioinformatics applications. Specifically, a 3-grams (sequence of three amino acids) is represented in the size of a 100-dimension vector to encode proteins for protein family classification (Asgari and Mofrad, 2015), which suggests state-of-the-art performance. To convert the influenza protein sequence into feature sets that can be managed by neural networks, we leverage ProtVec that can encode proteins through distributed representation for the construction of the proposed model.

To preserve the sequence information, the protein sequences of each segment are split into shifted overlapping residues in the window of 3 shown in Fig. 2. We take the HA segment as an example that each HA protein is 584 in length after alignment and is represented by 582 lists of 3-grams. Hence, each HA protein sequence is embedded in a 582*100 dimensional space based on ProtVec, where a 3-gram is denoted by a 100-dimension numerical vector. If a subsequence contains ‘-’ at any position, the ‘unknown’ vector will be assigned to represent the 3-grams. Similarly, the sequences of other influenza proteins are converted into (*N*-2)*100 dimensional vector that *N* stands for the length of distinct proteins.

**Fig. 2.**
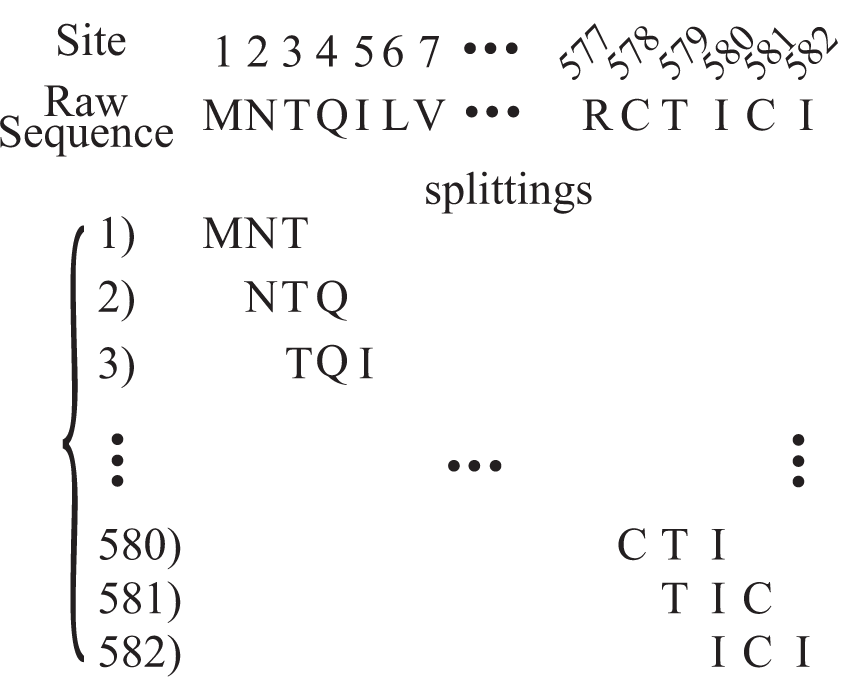
The splittings of influenza sequences after multiple sequence alignment. Each sequence is represented by 582 lists of 3-grams.

#### 2.3.2 CNN architecture

Convolutional neural network (CNN) has been successfully applied to a variety of fields for learning patterns from massive data, including image classification (Krizhevsky et al., 2012), action recognition (Wang et al., 2016a), protein secondary structure prediction (Wang et al., 2016b) and speech recognition (Huang et al., 2015). Ensemble learning has demonstrated its utility by generating multiple versions of a predictor network and using them to make aggregated predictions. The first revolutionary work using ensemble CNN was in computer vision for image classification tasks over the largest ImageNet database (Deng et al., 2009; Krizhevsky et al., 2012). After that, several other CNN architectures were proposed including VGG16, GoogleNet and ResNet (Simonyan and Zisserman, 2014; Szegedy et al., 2015; He et al., 2016), which exhibited very exciting performance in various problems. Besides, Chaib et al. improved regular VGG networks by fusing the output of the last two fully connected layers (Chaib et al., 2017). Roy et al. proposed a fused CNN for texture classification by merging the last representation layer of widely adapted AlexNet and VGG16 (Roy et al., 2020).

In this work, we leverage ResNeXt as the base model which demonstrates to achieve better performance than some similar CNN structures such as VGG and ResNet for virulence prediction on single segment dataset. It adopts highly modularized design following ResNets that consists of a stack of residual blocks and these blocks are subject to two rules inspired by ResNets with the same topology (Xie et al., 2017). The simplest neurons in artificial neural networks perform inner product and the output is the summation of *w*_*i*_ times *x*_*i*_. This can be recast as a combination of splitting, transformation and aggregating. The inner product can be represented as a form of aggregating transformation 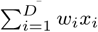, where *X* = [*x*_1_, *x*_2_, …, *x*_*D*_] is a D-channel input vector to the neuron and *w*_*i*_ is a filter’s weight for *i*-th channel. The elementary transformation (*w*_*i*_, *x*_*i*_) can be replaced with a more generic function after the analysis of simple neurons. Formally, the aggregated transformation is presented as:

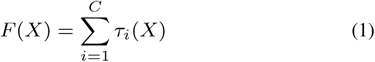

where *τ*_*i*_(*x*) can be an arbitrary function, *C* is the size of the set of transformations to be aggregated that is referred to cardinality. It is demonstrated that the accuracy can be gained more efficiently by increasing the cardinality than by going deeper or wider (Xie et al., 2017).

Fig. 3 shows a typical building block of ResNeXt. It is very similar to the Inception module (Szegedy et al., 2015) that they both follow the split-transform-merge paradigm, except in this variant, the outputs of different paths are merged by adding them together, while they are depth-concatenated in Inception model. In Eqn(1), a hyper-parameter called cardinality is introduced, which is the number of independent paths denoted as *C* to provide a new way of adjusting the capacity of the model. Compared to other structures such as Inception, the architecture of ResNeXt is easier to adapt to new datasets, as it has a simple paradigm and only one-hyperparameter to be adjusted. In addition, the aggregated transformation can be further served as the residue function formulated below, where *y* is the output.

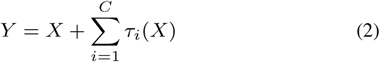

**Fig. 3.**
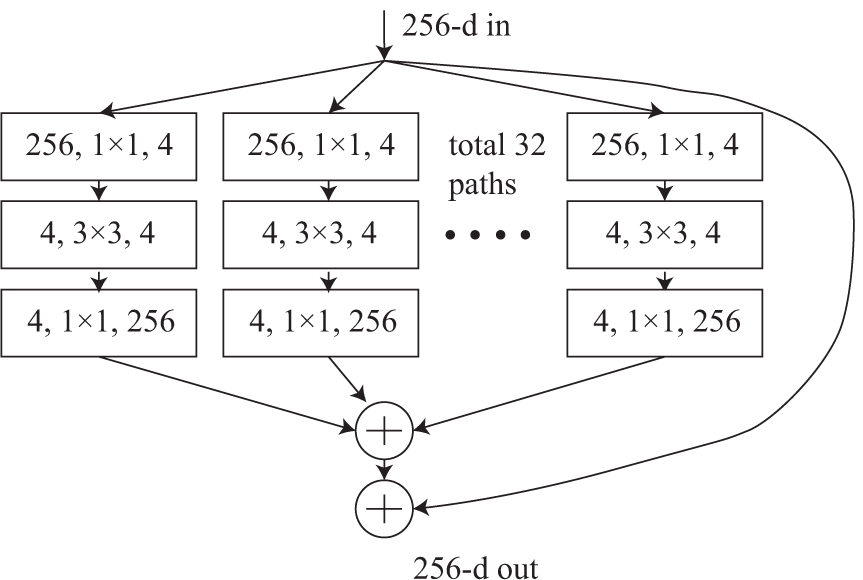
Aggregated residual transformation (Xie et al., 2017). A typical block of ResNeXt with cardinality = 32.

#### 2.3.3 Weighted ensemble CNN

To enhance the stability for the prediction of influenza virulence, a linear layer is introduced to combine the predictive results from all heads of the base ResNeXt model. Each head produces the prediction of virulence labels based on a single segment that partially suggests the degree of virulence. Assume *ya* is the predictive results from base model of VirPreNet, where *a* ∈ *S{*HA, NA, NP, PA, PB1, PB2, NS1, M1*}*. The final prediction can be formulated as:

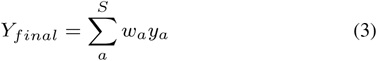

where *w*_*i*_ is the learned weight on the base model of segment *a* and *Y*_*final*_ is the final predictive outcome. Different weights will be assigned to the predictive results of the base ensemble model through the linear layer. By integrating the predictive results from all segments using base ensemble models, we can evaluate the predictions to find those that agree by decreasing the weight of inconsistent results to obtain a more reliable matching value. It has the ability to train on various datasets and concatenate their predictions to a final result that would help to achieve higher accuracy and capability for predictive tasks.

### 2.4 Experimental design

To evaluate the effectiveness of the proposed model, we test our model on the collected dataset with the MLD50-based definition of virulence. The experimental results indicate that VirPreNet obtains better predictive performance than the compared methods. Moreover, VirPreNet can also display the learned weights of base models built upon each segment for the final prediction. We will describe the experimental settings and provide a detailed performance comparison between the proposed model and baseline approaches.

#### 2.4.1 Baseline approaches

Three types of baselines are leveraged for influenza virulence prediction comparison. The first baseline directly applies three rule-based methods (Ivan and Kwoh, 2019) to detect the virulence of influenza A viruses. The second baseline is to use traditional machine learning approaches including logistic regression (LR), random forest (RF), neural network (NN) and K-nearest neighbor (KNN) for the predictive tasks. The third type of baseline comprises several CNN architectures including VGG-16, ResNet-34 and ResNeXt-50 for the comparison with VirPreNet using the same feature space. For all CNN-based models, the architecture is similar to the proposed model, but without a weighted mechanism using all 8 segments.

#### 2.4.2 Implementation

All the approaches are implemented with Rweka (Hornik et al., 2009), Scikit-learn (Pedregosa et al., 2011) and PyTorch (Paszke et al., 2017). The collected data is randomly divided into training and independent testing set in a ratio of 0.8:0.2. The 5-fold cross validation is applied in the training process while the testing set is used for validation on all approaches. We implement three rule-based methods for virulence prediction that are the same in (Ivan and Kwoh, 2019). For traditional machine learning methods, the parameters for these models are set by default. For all CNN-based models, we apply stochastic gradient descent with a minimum batch size of 4 for optimization. The learning rate is 0.001 and we set the size of filter window 10 with 130 filter maps. The L1 regularization and drop-out (rate = 0.5) strategy are carried out with 100 training epochs. The predictive performance is evaluated by accuracy, precision, recall and F-score for comparing the performance of all the models in virulence prediction.

## 3 Results and Discussion

In this experiment, we have proposed a model to learn distributed representation of amino acids from the influenza dataset. ResNeXt (Xie et al., 2017) is leveraged as our base model to build the VirPreNet framework. We first examine the effect of using different optimizers including Adaptive Moment Estimation (Adam) (Kingma and Ba, 2014), Adadelta (Zeiler, 2012), Adaptive Gradient Algorithm (AdaGrad) (Ruder, 2016), Root Mean Square Propagation (RMSProp) (Tieleman and Hinton, 2014) and Stochastic Gradient Descent (SGD) (Bottou, 2010) on our model. Next, we describe the ability of the ensemble CNN model on the virulence prediction over traditional approaches by exerting ProtVec. Finally, the comparative performance between VirPreNet and other individual segment-based ensemble CNN models is presented.

### 3.1 Optimizers ablation

Fig. 4 shows the predictive accuracy versus epochs with five different optimizers on the influenza dataset of the HA segment using the base model. The model is reinitialized after each round of optimization to provide a fair comparison between different optimizers. Overall, it can be observed that the model achieves a more stable result with over 0.65 in accuracy using the SGD optimizer. Though Adagrad optimizer indicates comparative prediction performance, it presents fierce fluctuation of predictive accuracy. The model obtains a more robust performance with Adadelta optimizer, whereas the average accuracy is lower than on the SGD optimizer. Similar patterns can be found when modeling on other single segment-based data (Supplementary Materials S1). Therefore, we have decided to choose SGD, an optimizer with superior performance to create our final model.

**Fig. 4.**
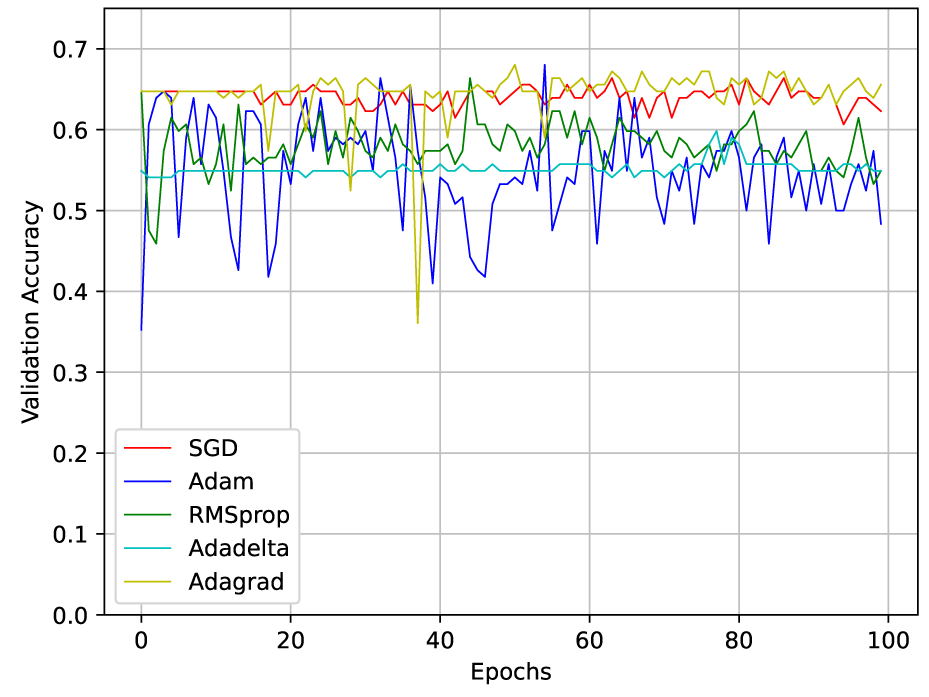
The validation accuracy on virulence prediction of influenza viruses using different optimizers on dataset of HA segment.

### 3.2 Comparative performance between ensemble CNN and traditional approaches on segment-based dataset

We examine the performance of using three types of baseline methods for virulence prediction on individual protein dataset. Table 1 shows the predictive results on HA dataset. It is shown that traditional machine learning classifiers outperform the rule-based methods on ProtVec-based features in terms of most measures. Moreover, deep learning-based models obtain competitive results compared with traditional machine learning approaches. ResNeXt-50 achieves the highest performance that is 0.676, 0.719 and 0.814 in accuracy, precision and F-score, respectively. They are 10.0%, 7.2% and 11.8% higher than the logistic regression model, which exhibits the best results among all other baselines. Interestingly, the VGG-16 model presents much better recall value over other approaches, this probably VGG-16 architecture is more sensitive to the true positive samples. The prediction performance using influenza datasets of other segments can be found in Supplementary Material S2. We only show the comparison between traditional machine learning approaches and deep learning-based models, since the rule-based methods perform worst among all the baselines. The results further demonstrate the advantage of ensemble CNN models on the virulence prediction of the influenza virus, specifically the ResNeXt-50 architecture, which will be chosen as our base model to create the proposed framework.

**Table 1.** Comparative performance between base model of VirPreNet and three types of baselines on HA segment dataset for virulence prediction.

| Model |  | Accuracy | Precision | Recall | F-score |
| --- | --- | --- | --- | --- | --- |
| Rule-Based | OneR | 0.495 | 0.502 | 0.496 | 0.499 |
|  | JRIP | 0.467 | 0.472 | 0.466 | 0.468 |
|  | PART | 0.553 | 0.562 | 0.557 | 0.559 |
| Traditional Machine Learning | RF | 0.566 | 0.645 | 0.717 | 0.679 |
|  | LR | 0.576 | 0.647 | 0.758 | 0.696 |
|  | KNN | 0.523 | 0.617 | 0.650 | 0.627 |
|  | NN | 0.539 | 0.636 | 0.667 | 0.644 |
| Deep Learning | VGG-16 | 0.645 | 0.645 | **1.000** | 0.784 |
|  | ResNet-34 | 0.666 | 0.674 | 0.936 | 0.783 |
|  | ResNeXt-50 | **0.676** | **0.719** | 0.938 | **0.814** |

### 3.3 Comparative performance between VirPreNet and ensemble base model

In statistical prediction, independent testing is often utilized to examine the capability of the predictor in practical application. To demonstrate the effectiveness of our proposed model, we compared VirPreNet with the single segment-based ResNeXt model on the prediction of influenza virulence. The 5-fold cross validation is leveraged in the training process, while the independent testing data is used to evaluate the ability of our model to predict new sample data. The results are averaged over 10 random trials and the details are shown in Table 2. It can be seen that VirPreNet achieves a noticeable higher performance than compared models. In more detail, VirPreNet can obtain an average value of 0.794 in accuracy, which is 10.8% higher than the best result of the ResNeXt-PB2 model. Regarding other evaluation metrics, the results also indicate that VirPreNet outperforms other models on the collected influenza dataset. Interestingly, the predictive performance based on PB2 and HA models is higher than others, which is consistent with the outcome from (Ivan and Kwoh, 2019).

**Table 2.** Comparative results between VirPreNet and base ensemble model.

| Model | Accuracy | Precision | Recall | F-score |
| --- | --- | --- | --- | --- |
| ResNeXt-PB2 | 0.686<br>(0.029) | 0.701<br>(0.048) | 0.875<br>(0.151) | 0.778<br>(0.048) |
| ResNeXt-PB1 | 0.648<br>(0.052) | 0.666<br>(0.037) | 0.914<br>(0.035) | 0.770<br>(0.031) |
| ResNeXt-PA | 0.648<br>(0.050) | 0.676<br>(0.048) | 0.889<br>(0.063) | 0.768<br>(0.029) |
| ResNeXt-HA | 0.676<br>(0.028) | 0.719<br>(0.041) | 0.938<br>(0.114) | 0.814<br>(0.029) |
| ResNeXt-NP | 0.666<br>(0.015) | 0.681<br>(0.030) | 0.921<br>(0.062) | 0.783<br>(0.006) |
| ResNeXt-NA | 0.668<br>(0.017) | 0.668<br>(0.011) | **0.965**<br>(**0.011**) | 0.790<br>(0.011) |
| ResNeXt-M1 | 0.637<br>(0.029) | 0.679<br>(0.029) | 0.853<br>(0.158) | 0.756<br>(0.055) |
| ResNeXt-NS1 | 0.666<br>(0.012) | 0.669<br>(0.013) | 0.959<br>(0.028) | 0.787<br>(0.006) |
| VirPreNet | **0.794**<br>(**0.028**) | **0.781**<br>(**0.044**) | 0.948<br>(0.029) | **0.856**<br>(**0.025**) |

Moreover, we plot the ROC (Receiver Operating Characteristic) curves with a significant proportion of area under curve (AUC) of all compared methods from Table 2. The AUC values display the matching degree with the experiment data ranging from 0 to 1. As can be seen in Fig. 5, VirPreNet can accurately predict the influenza virulence and outperform other single segment-based ensemble CNN models at almost all thresholds. Overall, the outcomes further demonstrate that our proposed model can better predict influenza virulence on the selected dataset, as well as make comprehensive improvement compared with baselines. It may also be applicable to predict the virulence of a wide range of viruses and drive the development of personalized medicine for infectious diseases.

**Fig. 5.**
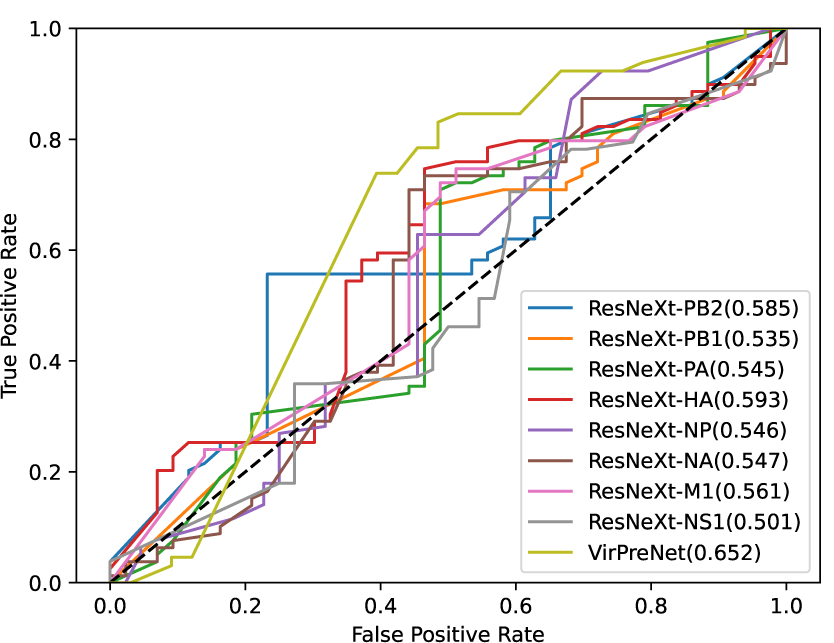
The ROC curves and AUC values for virulence prediction of all compared models generated from Table 2.

**Fig. 6.**
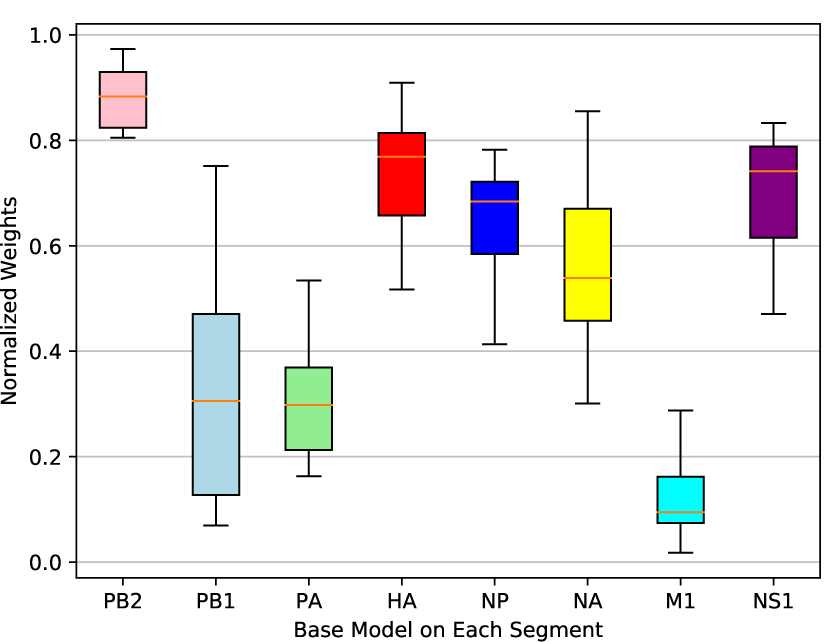
Learned weights by VirPreNet on the prediction of influenza virulence for the base ensemble models using single segment dataset. The X-axis represents the base models on the individual segment. As the values of the learned weights may be negative, the weights are normalized with softmax function. Y-axis represents the ranges of normalized weights by ten random trials.

### 3.4 Importance of single segment-based ensemble CNN model

Ensemble modeling implements the process where multiple diverse base models are incorporated to perform prediction tasks using different algorithms or datasets. In this work, the ensemble CNN models are utilized as the base model in our proposed model. The results suggest that VirPreNet improves the performance by aggregating the predication of each base model. To investigate the influence of each base model for the virulence prediction, we have visualized the learned weights shown in Fig. 5 for the contribution of individual models on the final prediction. These weights are obtained through the testing process from Table 2. Since the values of the learned weights may be negative, we use softmax function to normalize the weights. It can be observed that the PB2 segment contributes more to the final prediction outcome, followed by the HA segment. Interestingly, the models on other segments indicate less influence on virulence prediction. However, it does not mean that they are insignificant in the formation of virulence. One of the possible reasons is that we do not have sufficient sequence samples in the training process, which could greatly influence the predictive results.

The determination of influenza virulence has been investigated for decades, but it remains to be an open challenge, as the viral evolution is rapid and the mechanism of influenza strains to be virulent is very complicated and multi-factor associated. We utilized weight loss and survival data of infected mice to infer MLD50 values and converted them into classification tasks, for which our proposed method can be used for virulence prediction. However, for the evaluation of toxicity on humans, the best test species is human due to the interspecies variations between humans and other animals in physiology and biochemistry, in which the extrapolation of animal data may not be accurately applied to humans (Gallagher, 2002). Some clinical trials do not realize regulatory and marketing approval since it is not consistent in translating animal findings to the human situation (Shanks et al., 2009). Even though it remained controversial in transferring animal experiments to humans, it contributes a lot to our understanding of disease mechanisms and providing novel insights for further study (Van der Worp et al., 2010). For example, considering biomarkers on the influence of virulence, we can find mutations from different viral proteins. Graef et al. uncovered the PB2 subunit is a major virulence determinant of influenza viruses by interacting with the mitochondrial antiviral signaling protein and inhibiting expression of beta interferon (Graef et al., 2010). Yu et al. found adaptive changes that confer increased virulence in mammals on HA and PB2 segments using generated mouse-adapted viruses (Yu et al., 2018). Our proposed model has further revealed the importance of PB2 and HA segments on the inference of virulence compared with other segments through computational analysis. Nevertheless, more evidence is needed to identify what characteristics or mutations of PB2 or HA segment differ between virulent and avirulent influenza A strains.

## 4 Conclusion

In this paper, we propose a general weighted ensemble CNN framework named VirPreNet on virulence prediction of influenza A viruses using all 8 segments. Specifically, we employ MLD50 as criteria to label the virulence level of influenza strains. A continuous distributed representation is applied for the splittings and embeddings of genome sequences. ResNeXt is leveraged as the base model of VirPreNet to predict virulence based on the single viral segment. By integrating all the predictive results through a linear layer, the proposed framework can improve the performance of virulence prediction. We validate VirPreNet on the collected influenza dataset. Experimental results show that it outperforms existing machine learning methods and base ensemble CNN models. Moreover, the proposed model can automatically learn the weights for different base models. The analysis of the learned weights indicates the importance of the viral segment on the virulence prediction, which can provide novel insights into the formation of virulence and facilitate flu surveillance.

## 5 Supplementary Materials

The supplementary materials can be found: https://drive.google.com/open?id=1PB6402wbtrAeGIha3kBAt-S96AGwBzYS

## Acknowledgements

This project is supported by AcRF Tier 2 grant MOE2014-T2-2-023, Ministry of Education, Singapore and A*STAR-NTU-SUTD AI Partnership grant, RGANS1905.

## References

Altschul, S. F., Gish, W., Miller, W., Myers, E. W., and Lipman, D. J. (1990). Basic local alignment search tool. Journal of molecular biology, 215(3):403–410.

Asgari, E. and Mofrad, M. R. (2015). Continuous distributed representation of biological sequences for deep proteomics and genomics. PloS one, 10(11):e0141287.

Bottou, L. (2010). Large-scale machine learning with stochastic gradient descent. In Proceedings of COMPSTAT’2010, pages 177–186. Springer.

Bouvier, N. M. and Palese, P. (2008). The biology of influenza viruses. Vaccine, 26:D49–D53.

Burke, D. F. and Smith, D. J. (2014). A recommended numbering scheme for influenza a ha subtypes. PloS one, 9(11):e112302.

Chaib, S., Liu, H., Gu, Y., and Yao, H. (2017). Deep feature fusion for vhr remote sensing scene classification. IEEE Transactions on Geoscience and Remote Sensing, 55(8):4775–4784.

Cheung, P. P.-H., Rogozin, I. B., Choy, K.-T., Ng, H. Y., Peiris, J. S. M., and Yen, H.-L. (2015). Comparative mutational analyses of influenza a viruses. Rna, 21(1):36–47.

Chou, K.-C. (2015). Impacts of bioinformatics to medicinal chemistry. Medicinal chemistry, 11(3):218–234.

da Costa, L. M., Grella, T. C., Barbosa, R. A., Malaspina, O., and Nocelli, R. C. F. (2015). Determination of acute lethal doses (ld50 and lc50) of imidacloprid for the native bee melipona scutellaris latreille, 1811 (hymenoptera: Apidae). Sociobiology, 62(4):578–582.

de Vries, E., de Vries, R. P., Wienholts, M. J., Floris, C. E., Jacobs, M.-S., van den Heuvel, A., Rottier, P. J., and de Haan, C. A. (2012). Influenza a virus entry into cells lacking sialylated n-glycans. Proceedings of the National Academy of Sciences, 109(19):7457–7462.

Deng, J., Dong, W., Socher, R., Li, L.-J., Li, K., and Fei-Fei, L. (2009). Imagenet: A large-scale hierarchical image database. In 2009 IEEE conference on computer vision and pattern recognition, pages 248–255. Ieee.

Gallagher, M. E. (2002). Toxicity testing requirements, methods and proposed alternatives. Environs: Envtl. L. & Pol’y J., 26:253.

Garg, A. and Gupta, D. (2008). Virulentpred: a svm based prediction method for virulent proteins in bacterial pathogens. BMC bioinformatics, 9(1):62.

Graef, K. M., Vreede, F. T., Lau, Y.-F., McCall, A. W., Carr, S. M., Subbarao, K., and Fodor, E. (2010). The pb2 subunit of the influenza virus rna polymerase affects virulence by interacting with the mitochondrial antiviral signaling protein and inhibiting expression of beta interferon. Journal of virology, 84(17):8433–8445.

He, K., Zhang, X., Ren, S., and Sun, J. (2016). Deep residual learning for image recognition. In Proceedings of the IEEE conference on computer vision and pattern recognition, pages 770–778.

Hornik, K., Buchta, C., and Zeileis, A. (2009). Open-source machine learning: R meets weka. Computational Statistics, 24(2):225–232.

Huang, J.-T., Li, J., and Gong, Y. (2015). An analysis of convolutional neural networks for speech recognition. In 2015 IEEE International Conference on Acoustics, Speech and Signal Processing (ICASSP), pages 4989–4993. IEEE.

Imai, M. and Kawaoka, Y. (2012). The role of receptor binding specificity in interspecies transmission of influenza viruses. Current opinion in virology, 2(2):160–167.

Ivan, F. X. and Kwoh, C.-K. (2019). Rule-based meta-analysis reveals the major role of pb2 in influencing influenza a virus virulence in mice. bioRxiv, page 556647.

Kamal, R. P., Katz, J. M., and York, I. A. (2014). Molecular determinants of influenza virus pathogenesis in mice. In Influenza Pathogenesis and Control-Volume I, pages 243–274. Springer.

Katoh, K. and Standley, D. M. (2013). Mafft multiple sequence alignment software version 7: improvements in performance and usability. Molecular biology and evolution, 30(4):772–780.

Kingma, D. P. and Ba, J. (2014). Adam: A method for stochastic optimization. arXiv preprint 1412.6980.

Krizhevsky, A., Sutskever, I., and Hinton, G. E. (2012). Imagenet classification with deep convolutional neural networks. In Advances in neural information processing systems, pages 1097–1105.

LeGoff, J., Rousset, D., Abou-Jaoude, G., Scemla, A., Ribaud, P., Mercier-Delarue, S., Caro, V., Enouf, V., Simon, F., Molina, J.-M., et al. (2012). I223r mutation in influenza a (h1n1) pdm09 neuraminidase confers reduced susceptibility to oseltamivir and zanamivir and enhanced resistance with h275y. PLoS One, 7(8):e37095.

Ma, M.-J., Liu, C., Wu, M.-N., Zhao, T., Wang, G.-L., Yang, Y., Gu, H.-J., Cui, P.-W., Pang, Y.-Y., Tan, Y.-Y., et al. (2018). Influenza a (h7n9) virus antibody responses in survivors 1 year after infection, china, 2017. Emerging infectious diseases, 24(4):663.

Ndifon, W., Dushoff, J., and Levin, S. A. (2009). On the use of hemagglutination-inhibition for influenza surveillance: surveillance data are predictive of influenza vaccine effectiveness. Vaccine, 27(18):2447–2452.

Organization, W. H. et al. (2009). Fact sheet no 211: Influenza (seasonal). WHO: Geneva, Switzerland, April.

Paszke, A., Gross, S., Chintala, S., Chanan, G., Yang, E., DeVito, Z., Lin, Z., Desmaison, A., Antiga, L., and Lerer, A. (2017). Automatic differentiation in pytorch.

Pedregosa, F., Varoquaux, G., Gramfort, A., Michel, V., Thirion, B., Grisel, O., Blondel, M., Prettenhofer, P., Weiss, R., Dubourg, V., et al. (2011). Scikit-learn: Machine learning in python. Journal of machine learning research, 12(Oct):2825–2830.

Pirofski, L.-a. and Casadevall, A. (2012). Q&a: What is a pathogen? a question that begs the point. BMC biology, 10(1):6.

Poovorawan, Y., Pyungporn, S., Prachayangprecha, S., and Makkoch, J. (2013). Global alert to avian influenza virus infection: from h5n1 to h7n9. Pathogens and global health, 107(5):217–223.

Roy, S. K., Dubey, S. R., Chanda, B., Chaudhuri, B. B., and Ghosh, D. K. (2020). Texfusionnet: An ensemble of deep cnn feature for texture classification. In Proceedings of 3rd International Conference on Computer Vision and Image Processing, pages 271–283. Springer.

Ruder, S. (2016). An overview of gradient descent optimization algorithms. arXiv preprint 1609.04747.

Saunders-Hastings, P. R. and Krewski, D. (2016). Reviewing the history of pandemic influenza: understanding patterns of emergence and transmission. Pathogens, 5(4):66.

Sayers, E. W., Barrett, T., Benson, D. A., Bolton, E., Bryant, S. H., Canese, K., Chetvernin, V., Church, D. M., DiCuccio, M., Federhen, S., et al. (2012). Database resources of the national center for biotechnology information. Nucleic acids research, 40(D1):D13–D25.

Seo, S. H., Hoffmann, E., and Webster, R. G. (2002). Lethal h5n1 influenza viruses escape host anti-viral cytokine responses. Nature medicine, 8(9):950.

Shanks, N., Greek, R., and Greek, J. (2009). Are animal models predictive for humans? Philosophy, ethics, and humanities in medicine, 4(1):2.

Simonyan, K. and Zisserman, A. (2014). Very deep convolutional networks for large-scale image recognition. arXiv preprint 1409.1556.

Song, J., Xu, J., Shi, J., Li, Y., and Chen, H. (2015). Synergistic effect of s224p and n383d substitutions in the pa of h5n1 avian influenza virus contributes to mammalian adaptation. Scientific reports, 5:10510.

Su, C. T.-T., Ouyang, X., Zheng, J., and Kwoh, C.-K. (2013). Structural analysis of the novel influenza a (h7n9) viral neuraminidase interactions with current approved neuraminidase inhibitors oseltamivir, zanamivir, and peramivir in the presence of mutation r289k. BMC bioinformatics, 14(16):S7.

Su, S., Bi, Y., Wong, G., Gray, G. C., Gao, G. F., and Li, S. (2015). Epidemiology, evolution, and recent outbreaks of avian influenza virus in china. Journal of virology, 89(17):8671–8676.

Szegedy, C., Liu, W., Jia, Y., Sermanet, P., Reed, S., Anguelov, D., Erhan, D., Vanhoucke, V., and Rabinovich, A. (2015). Going deeper with convolutions. In Proceedings of the IEEE conference on computer vision and pattern recognition, pages 1–9.

Thrall, P. H. and Burdon, J. J. (2003). Evolution of virulence in a plant host-pathogen metapopulation. Science, 299(5613):1735–1737.

Tieleman, T. and Hinton, G. (2014). Rmsprop gradient optimization. URL http://www.cs.toronto.edu/tijmen/csc321/slides/lecture_slides_lec6.pdf.

Van der Worp, H. B., Howells, D. W., Sena, E. S., Porritt, M. J., Rewell, S., O’Collins, V., and Macleod, M. R. (2010). Can animal models of disease reliably inform human studies? PLoS med, 7(3):e1000245.

Vijaykrishna, D., Mukerji, R., and Smith, G. J. (2015). Rna virus reassortment: an evolutionary mechanism for host jumps and immune evasion. PLoS pathogens, 11(7):e1004902.

Wang, P., Cao, Y., Shen, C., Liu, L., and Shen, H. T. (2016a). Temporal pyramid pooling-based convolutional neural network for action recognition. IEEE Transactions on Circuits and Systems for Video Technology, 27(12):2613–2622.

Wang, S., Peng, J., Ma, J., and Xu, J. (2016b). Protein secondary structure prediction using deep convolutional neural fields. Scientific reports, 6:18962.

Wu, Y., Cho, M., Shore, D., Song, M., Choi, J., Jiang, T., Qi, J., Li, A., Yi, K. S., Chang, M., et al. (2016). Micro neutralization (mn) assay of influenza viruses with monoclonal antibodies.

Xie, S., Girshick, R., Dollár, P., Tu, Z., and He, K. (2017). Aggregated residual transformations for deep neural networks. In Proceedings of the IEEE conference on computer vision and pattern recognition, pages 1492–1500.

Yamada, S., Hatta, M., Staker, B. L., Watanabe, S., Imai, M., Shinya, K., Sakai-Tagawa, Y., Ito, M., Ozawa, M., Watanabe, T., et al. (2010). Biological and structural characterization of a host-adapting amino acid in influenza virus. PLoS pathogens, 6(8):e1001034.

Yin, R., Luusua, E., Dabrowski, J., Zhang, Y., and Kwoh, C. K. (2020a). Tempel: time-series mutation prediction of influenza a viruses via attention-based recurrent neural networks. Bioinformatics.

Yin, R., Zhou, X., Ivan, F. X., Zheng, J., Chow, V. T., and Kwoh, C. K. (2017). Identification of potential critical virulent sites based on hemagglutinin of influenza a virus in past pandemic strains. In Proceedings of the 6th International Conference on Bioinformatics and Biomedical Science, pages 30–36. ACM.

Yin, R., Zhou, X., Rashid, S., and Kwoh, C. K. (2020b). Hopper: an adaptive model for probability estimation of influenza reassortment through host prediction. BMC Medical Genomics, 13(1):9.

Yu, Z., Cheng, K., Sun, W., Zhang, X., Xia, X., and Gao, Y. (2018). Pb2 and ha mutations increase the virulence of highly pathogenic h5n5 clade 2.3. 4.4 avian influenza virus in mice. Archives of virology, 163(2):401–410.

Zeiler, M. D. (2012). Adadelta: an adaptive learning rate method. arXiv preprint 1212.5701.

Zhang, W., Shi, Y., Lu, X., Shu, Y., Qi, J., and Gao, G. F. (2013). An airborne transmissible avian influenza h5 hemagglutinin seen at the atomic level. Science, 340(6139):1463–1467.

Zheng, L.-L., Li, Y.-X., Ding, J., Guo, X.-K., Feng, K.-Y., Wang, Y.-J., Hu, L.-L., Cai, Y.-D., Hao, P., and Chou, K.-C. (2012). A comparison of computational methods for identifying virulence factors. PLoS One, 7(8).

Zhou, X., Zheng, J., Ivan, F. X., Yin, R., Ranganathan, S., Chow, V. T., and Kwoh, C.-K. (2018). Computational analysis of the receptor binding specificity of novel influenza a/h7n9 viruses. BMC genomics, 19(2):88.

